# IFNγ initiates TLR9-dependent autoimmune hepatitis in DNase II deficient mice

**DOI:** 10.1101/2024.07.10.602775

**Authors:** Kaiyuan Hao, Kevin MingJie Gao, Melissa Strauss, Sharon Subramanian, Ann Marshak-Rothstein

## Abstract

Patients with biallelic hypomorphic mutation in *DNASE2* develop systemic autoinflammation and early-onset liver fibrosis. Prior studies showed that Dnase2^-/-^ Ifnar^-/-^ double knockout (DKO) mice develop Type I IFN-independent liver inflammation, but immune mechanisms were unclear. We now show that DKO mice recapitulate many features of human autoimmune hepatitis (AIH), including periportal and interstitial inflammation and fibrosis and elevated ALT. Infiltrating cells include CD8+ tissue resident memory T cells, type I innate lymphoid cells, and inflammatory monocyte/macrophage cells that replace the Kupffer cell pool. Importantly, TLR9 expression by bone marrow-derived cells is required for the the development of AIH. TLR9 is highly expressed by inflammatory myeloid cells but not long-lived Kupffer cells. Furthermore, the initial recruitment of TLR9 expressing monocytes and subsequent activation of lymphocytes requires IFNγ signaling. These findings highlight a critical role of feed forward loop between TLR9 expressing monocyte-lineage cells and IFNg producing lymphocytes in autoimmune hepatitis.

## INTRODUCTION

Failure to appropriately clear DNA from extracellular, cytosolic, or endosomal/lysosomal compartments can lead to a spectrum of debilitating autoimmune or autoinflammatory diseases driven by the aberrant activation of DNA pattern recognition receptors. For example, loss of the extracellular DNAse, DNAse1L1, can lead to activation of endosomal TLRs and the ensuing development of SLE (1), while hypomorphic mutations in the cytosolic DNAse, Trex1, lead to the activation of the cGAS/STING pathway and causes Aicardi-Goutieres Syndrome (**AGS**) (2, 3). Many of these conditions have been designated monoclonal type I interferonopathies, since these patients have a strong IFN signature, as defined by the upregulation of IFN-stimulated genes (**ISGs**) (4, 5). However, IFN signatures may be attributable to either type I or type II (IFN*γ*) responses and IFN*γ* is well recognized as a critical enhancer of macrophage innate immune responses (6). Importantly, both endosomal and cytosolic NA sensors have been shown to induce both type I and type II IFN responses. To better understand the interplay between NA sensors, IFNs, and the effector cells responsible for the pathogenesis of these autoinflammatory conditions, animal models that recapitulate the human disease syndromes can be used to explore causal molecular mechanisms.

Patients with biallelic hypomorphic mutations in the endolysosomal DNAse, *DNase2,* develop an autoinflammatory disease associated with a range of clinical manifestations that impact the joints, skin, brain and liver (7). DNase II is a lysosomal endonuclease responsible for the degradation of DNA derived from phagocytosed apoptotic cells or other cell debris. Loss of function in DNase II results in excessive accumulation of undigested DNA in phagolysosomes, where it can activate endosomal TLRs, or in the cytosol where it can engage cytosolic NA sensors, thereby leading to immune abnormalities (8). DNase II deficiency in mice causes severe anemia and embryonic lethality that can be rescued by intercrossing with IFN*a*R-deficient or STING-deficient mice (9, 10). Although they lack type I interferon signaling, *Dnase2^-/-^ Ifnar^-/-^* double knockout (**DKO**) mice nevertheless develop autoimmune phenotypes that recapitulate clinical manifestations of *DNASE2* hypomorphic patients. The key features include a well-studied late-onset polyarthritis that depends on the cGAS/STING pathway and the absent in melanoma 2 (**AIM2**) inflammasome (8, 11–13). In addition, from early life, DKO mice exhibit symptoms of systemic autoimmunity such as autoantibody production, splenomegaly, elevated serum cytokines, and activated lymphocytes, all of which depend on expression of functional Unc93B1 and thus endosomal TLR signaling (11, 14). DKO mice also present with autoimmune hepatitis (**AIH**), also described in the *DNASE2* hypomorphic patients. Existing murine models of AIH are very limited (15–17),. Therefore,DKO mice provide a unique opportunity for both evaluating the role of nucleic acid sensors and IFN*γ* in systemic autoinflammatory diseases and also for identifying the specific cell types and mechanisms responsible for AIH. The current report defines the specific liver-infiltrating inflammatory subsets associated with DKO hepatitis and identifies a key role for both TLR9 and IFN*γ* in DKO liver pathology.

## RESULTS

### DKO mice develop an inflammatory liver disease associated with infiltrating myeloid cells, ILC1 cells, and CD8^+^ T cells

#### Clinical manifestations of hepatitis in DKO mice

The early onset inflammatory liver disease that develops in DKO mice recapitulates many of the clinical features of type 1 AIH as described in patient populations (18). These include severe portal mononuclear inflammation, extensive periportal and interstitial fibrosis, interface hepatitis as reflected by increased hepatocyte cell death at the inflammatory border (**Fig 1A**), the production of anti-nuclear antibodies (**ANA**) (19), and elevated serum levels of the liver enzyme ALT, indicative of hepatocyte damage (**Fig 1B**). It was important to better characterize the cell types contributing to the inflammatory infiltrate in order to identify potential effector mechanisms and also to establish criteria for comparing additional gene-targeted strains, as described below. Cell suspensions obtained from the livers of DKO and *Dnase^+/-^ Ifnar^-/-^* (**Het**) littermate controls were compared by flow cytometry. The total number of CD45^+^ hematopoietic cells isolated from DKO livers was increased by over 4-fold compared to the Het mice, with particularly increased numbers of CD11b^+^ myeloid cells, NK1.1^+^ cells, and CD8^+^ T cells (**Fig. 1C**).

**Figure 1.**
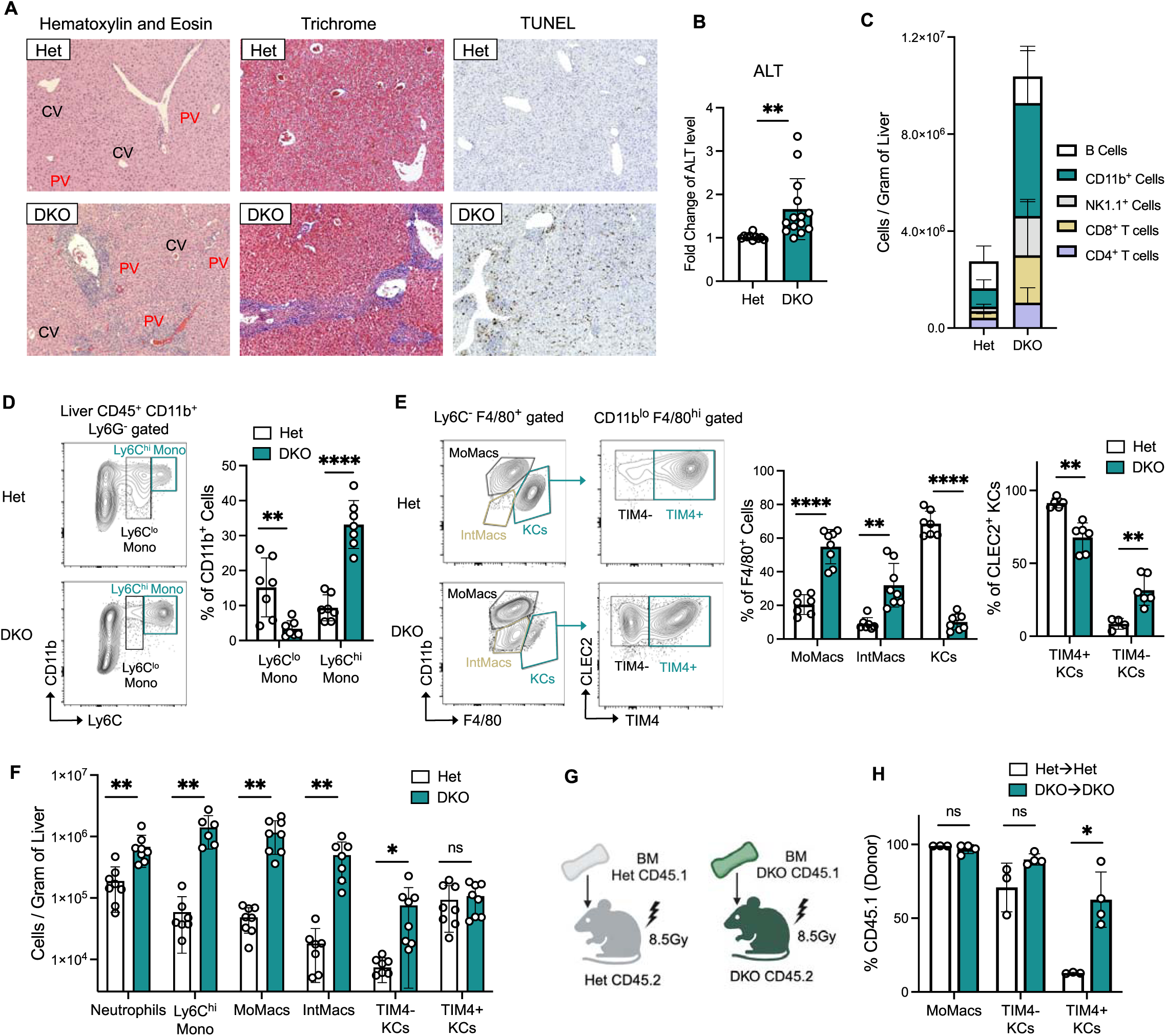
Infiltration of different myeloid subsets in a DNase II deficiency-induced autoimmune hepatitis mouse model. (A) Representative H&E, Masson’s trichrome, and TUNEL staining (10x) of liver sections from 3-4 month-old mice. (B) Relative ALT levels in the sera of 3-4 month-old mice normalized to Het controls (n = 4-8 per group). (C) Absolute numbers of indicated immune subsets within liver CD45^+^ immune cells (n = 6-15 per group). (D) Representative Ly6C and CD11b staining of CD11b^+^ Ly6G^-^ gated liver immune cells of 3-4 month-old mice. Bar graph compares the frequency of Ly6C^hi^ monocytes and Ly6C^lo^ monocytes within CD11b^+^ liver myeloid cells (n = 7 per group). (E) Representative CD11b and F4/80 staining of CD11b^+^ Ly6G^-^ Ly6C^-^ F4/80^+^ gated liver macrophages and CLEC2 and TIM4 staining of CD11b^+^ Ly6G^-^ Ly6C^-^ F4/80^hi^ CD11b^lo^ (KCs) gated cells of 3-4 month-old mice. Bar graphs compare the frequency of each macrophage subset within liver F4/80+ cells and each KC subset within F4/80^hi^ CD11b^lo^ cells (n = 6-7 per group). (F) Absolute cell counts of indicated myeloid subsets in the livers of 3-4 month-old mice (n = 6-8 per group). (G) Generating bone marrow chimeras using CD45.1 mice as donors and CD45.2 mice as hosts. (H) Extent of chimerism for different macrophage subsets 8 weeks post bone marrow transplant (n = 3-4 per group). Data are pooled from at least 2 independent experiments and represented as mean ± SEM. *p < 0.05, **p < 0.01, ***p < 0.001, and ****p < 0.0001. See also Figure S1.

#### Myeloid cells

CD11b^+^ myeloid cells constituted ∼40% of the total number of CD45^+^ cells in the DKO livers, and the vast majority of DKO infiltrating myeloid cells could be considered inflammatory myeloid lineage subsets. These included CD11b^+^ Ly6C^hi^ inflammatory macrophages, CD11b^hi^ F4/80^+^ monocyte-derived macrophages (MoMacs) and CD11b^lo^ F4/80^+^ intermediate macrophages (Int-Macs) (**Figure 1D,E,F**). By contrast, the Het livers contained a much lower percentage of these inflammatory subsets. Instead, the majority of Het myeloid cells were F480^hi^ macrophages, that included both mature macrophages and Kupffer cells (**KC**) (**Figure 1E,F**). KCs are normally yolk sac-derived self-renewing tissue resident macrophages that line the liver sinusoids and play a critical role in the detection and removal of bacteria and cell debris. KCs can also produce immune mediators such as inflammatory cytokines and oxygen radicals (20). They are routinely identified as CD11b^lo^, F4/80^hi^, CLEC2^+^, TIM4^+^ cells by flow cytometry. We also observed more CD11b^+^ Ly6C^lo^ patrolling monocytes in the Het livers (**Figure 1D**). Patrolling monocytes survey the apical side of the endothelium, sensing and removing damaged endothelial cells or other harmful particles (21).

KC constituted a much smaller proportion of the total myeloid population in DKO livers and included both a long-lived TIM4^+^ fraction as well as a TIM4^-^ subset, most likely derived from infiltrating monocytes (22). To compare the origin of the CLEC2^+^ KCs in WT and DKO mice, DKO and Het bone marrow (**BM**) stem cells were used to reconstitute lethally irradiated CD45-allelically distinct DKO and Het recipients, respectively, to generate DKO CD45.1àDKO CD45.2 and Het CD45.1àHet CD45.2 radiation chimeras. 8 weeks later, CD45 allelic markers were used to track the stem cell contribution to the myeloid lineage subsets (**Figure 1G,H**). As expected, in the HetàHet chimeras, all the inflammatory monocytes, monocyte-derived macrophages, and the majority of the TIM4^-^ KCs were derived from BM stem cells, while most of the TIM4^+^ KC remained host derived (**Figure 1H**). By contrast, in the DKOàDKO chimeras, not only were the inflammatory monocytes, monocyte-derived macrophages and TIM4^-^ KC derived from the BM stem cells, but the majority of the TIM4^+^ KC were also donor-derived. These data indicate that tissue resident KC in Het mice are long-lived and even survive lethal irradiation. However, the survival of long-lived TIM4^+^ KCs is compromised in DKO mice where this subset is then replaced with newly differentiated KC derived from stem cell derived infiltrating monocytes. To investigate the proliferative status of these cell types in Het and DKO mice, we measured expression levels of Ki-67 and a marker of cellular proliferation. Monocytes in both Het and DKO mice were nearly universally Ki-67 positive, reflecting an actively proliferating population (**Supplemental Figure 1A**). By contrast, the vast majority of TIM4^+^ KCs are negative for Ki-67, reflecting a more long-lived and quiescent phenotype (23), while TIM4^-^ KCs, showed an intermediate Ki-67 phenotype. Consistently, the proportion and total number of MoMacs and TIM4^-^ KCs were elevated in DKO liver (Figure 1E,F), These data are consistent with the notion that inflammatory monocytes proliferate in the liver and differentiate into more mature macrophage subsets(23), although we cannot rule out the possibility of self-renewing KCs, especially in the Het mice.

#### NK1.1^+^ lymphocytes

The proportion and the total number of NK1.1^+^ cells were also dramatically increased in the DKO mice as compared to the Het controls. The NK1.1^+^ marker identifies two major types of innate lymphocytes, conventional NK1.1^+^ DX5^+^ natural killer cells (**cNK**), and NK1.1^+^ CD49^+^ type I innate lymphoid cells (**ILC1**) (24). In the Het liver, the relatively small number of NK1.1^+^ cells were mainly cNK cells or double negative immature NK precursors (25) (**Figure 2A**). By contrast, there were very few cNK cells in the DKO liver. Instead, the NK1.1^+^ cells in the DKO mice were mainly ILC1 cells. Thus, the expanded subset of ILC1 is a distinct feature of DKO mice.

**Figure 2.**
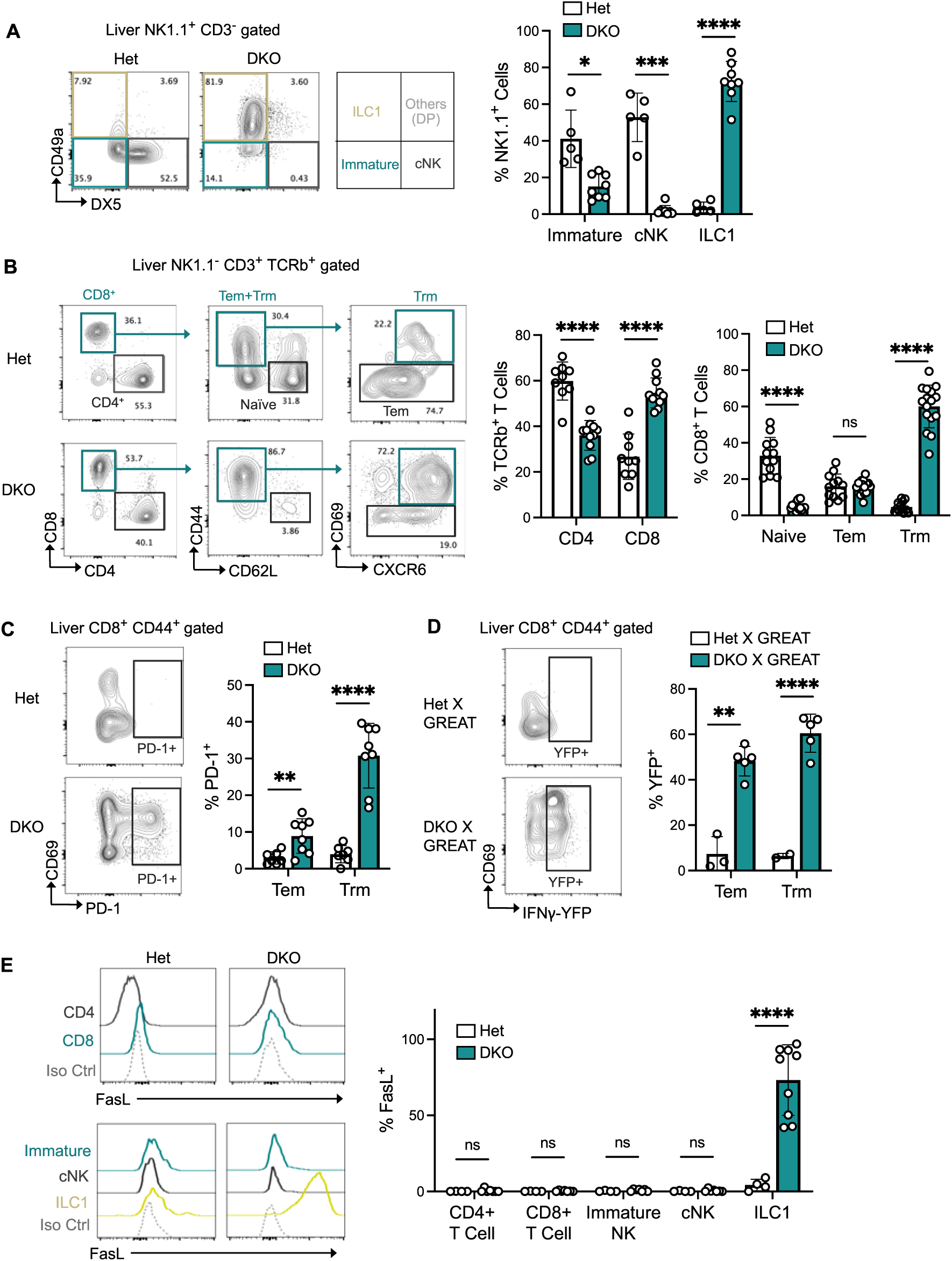
Liver infiltrating lymphocytes in DKO mice remain activated and acquire distinct effector functions. (A) Representative CD49a and DX5 staining of NK1.1^+^ CD3^-^ gated liver immune cells of 3-4 month-old mice. Bar graph compares the frequency of Type I innate lymphoid cells (ILC1), immature NK cells, and conventional NK cells (cNK) within liver NK1.1^+^ cells (n = 5-8 per group). (B) Gating strategy to identify CD8^+^ naïve T cells, effector memory T cells (Tem), and tissue-resident memory T cells (Trm) in livers of 3-4 month-old mice. Bar graphs compare the frequency of CD4^+^ and CD8^+^ T cells within liver T cell population, and frequency of naïve, Tem, and Trm cells within CD8^+^ T cells (n = 9-16 per group). (C) Representative CD69 and PD-1 staining of CD44^+^ gated liver CD8^+^ T cells of 3-4 month-old mice. Bar graph compares the frequency of PD-1^+^ exhausted T cells within CD8^+^ Tem and Trm cells respectively (n = 6-8 per group). (D) Representative CD69 staining and YFP flow plot of CD44^+^ gated activated liver CD8^+^ T cells of 3-4 month-old mice. Bar graph compares the frequency of YFP+ cells within CD8^+^ Tem and Trm cells separately (n = 3-5 per group). (E) Histogram of cell surface FasL expression on indicated cell types in the livers of 3-4 month-old mice (n = 4-9 per group). Data are pooled from at least 2 independent experiments and represented as mean ± SEM. *p < 0.05, **p < 0.01, ***p < 0.001, and ****p < 0.0001. See also Figure S2.

#### CD8^+^ T resident memory subsets

The ratio of CD8^+^ to CD4^+^ T cells shifted significantly in the DKO mice due to a marked expansion in the number of the CD8^+^ cells (**Figure 2B**). In the absence of any obvious infectious, the level of expansion points to an antigen-driven response, most likely directed at self-antigens. Not only were the number of CD8^+^ cells increased, but they were significantly more activated as indicated by loss of CD62L and high expression of CD44 (**Figure 2B**). In the context of infection, the combination of antigen, co-stimulation and inflammation commonly leads to the differentiation of relatively short-lived CXCR3+ cytotoxic effector cells that die off once the antigen is cleared. Activated T cells can also persist as a heterogeneous pool of memory subsets that have been broadly categorized as T effector memory (**Tem**) and T resident memory (**Trm**) as distinguished by the expression of markers such as CD69 and CXCR6 (**Figure 2B**). Tem cells are thought to produce IFN*γ*, survey peripheral sites, and retain the capacity to rapidly acquire cytotoxic activity if they reencounter antigen. Trm cells are defined as a more long-lived peripheral tissue-resident subset and may include chronically stimulated and exhausted T cell subsets (**Tex**). Tex cells are thought to have more limited effector activity and are identified by the expression of a variety of inhibitory receptors such as PD-1. Importantly, most of the activated cells in the Het mice were Tem cells. By contrast, most of the CD8^+^ T cells in the DKO mice were activated Trm cells. The proportion of naïve CD8^+^ T cells was much higher in the Het liver while the proportion of Trm was much higher in the DKO liver (**Figure 2B**). In addition, the DKO Trm subset contained a considerable number of cells that expressed PD-1, an indication of prolonged stimulation and at least partial exhaustion (**Figure 2C**). As a metric of CD8^+^ T cell function, we used the IFN*γ*-reporter line, GREAT (26), to assess IFN*γ* production. Intriguingly, a remarkably high frequency of CD8^+^ Trm cells are constitutively producing IFN*γ* (**Figure 2D**). Overall, these data indicate that autoinflammation in the DKO liver results in chronic self-antigen driven activation of CD8^+^ T cells, and that CD8^+^ T effector cells are likely participants in hepatic autoinflammation.

#### CD4^+^ T resident subsets

Although CD4^+^ cells in the liver were not as dramatically expanded as far as number, CD4^+^ T cells in the DKO mice were also more activated than T cells in the Het mice. In addition, significantly more CD4^+^ T cells in the DKO mice expressed PD-1 and made IFN*γ*, when compared to the Het control groups (**Supplemental Figure 2**). Therefore, chronically activated CD4^+^ T cells likely also accumulate in DKO livers.

#### FasL is highly expressed by liver infiltrating ILC1 cells

T cells express a range of molecules that can damage or injure cells through direct cytotoxicity and/or by promoting inflammation. One example is FasL, often expressed by cytotoxic cells, and in particular, Th1 cells that also produce IFN*γ*. Fas/FasL interactions are highly relevant to the liver, as now classic studies have shown that mice injected i.v. with anti-Fas antibody die as a result of severe liver damage (27–29). Moreover, hepatocytes constitutively express Fas and FasL-expressing cells have been shown to directly induce hepatocyte apoptosis (27). FasL has also been implicated in hepatitis and LPS-induced mortality (30). To determine whether any of the T cells that accumulate in the liver of DKO mice express FasL, liver cell suspensions from DKO, Het, and FasL-deficient mice were co-stained with lineage-specific markers and an anti-FasL antibody, MFL3. Unexpectedly, neither CD4^+^ nor CD8^+^ T cells in the livers of DKO mice expressed detectable levels of FasL (**Figure 2E)**. However, FasL was very highly expressed by ILC1 cells in DKO livers but not by any NK1.1^+^ cells in the Het controls. We compared the signal intensity to FasL-APC isotype control to ensure the signal is not due to non-specific binding. These data implicate ILC1 cells as directly contributing to hepatocyte death in DKO mice.

### Role of endosomal TLRs in systemic and hepatic inflammation of DKO mice

In the absence if DNAse, excessive DNA accumulates in phagolysosomal compartments, and therefore we initially assumed that the Unc93B-dependent DNA sensor, TLR9, would be the pattern recognition receptor responsible for lupus-related immune abnormalities in DKO mice. One of the distinguishing features of patients with DNase2 hypomorphic mutations is the development of lupus-like autoantibodies, and it is now well established that autoantibody production in SLE depends on B cell expression of endosomal TLRs (31). In fact, we have found that DKO mice also make ANAs through an Unc93B1-dependent mechanism, but, unexpectedly, the DKO sera gave speckled Hep-2 immunofluorescent staining patterns normally found in TLR7-driven models of SLE (7, 14), and not a homogeneous nuclear/mitotic plate staining pattern typical of anti-dsDNA antibodies. Therefore, we assumed that cell debris, resulting from the death of oversaturated DKO scavenger cells, provided the ligands that activated autoreactive DKO B cells specific for RNA-binding proteins, thereby leading to the production of autoantibodies and triggering other systemic manifestations of autoimmunity in DKO mice. To test this hypothesis, we generated TLR7-deficient TKO mice, and as expected, they did not make autoantibodies (**Figure 3A**). However, they still developed splenomegaly and hepatitis (**Figures 3B,C**). These results pointed to a nucleic acid-sensor other than TLR7 in DKO systemic autoinflammation and liver pathology.

**Figure 3.**
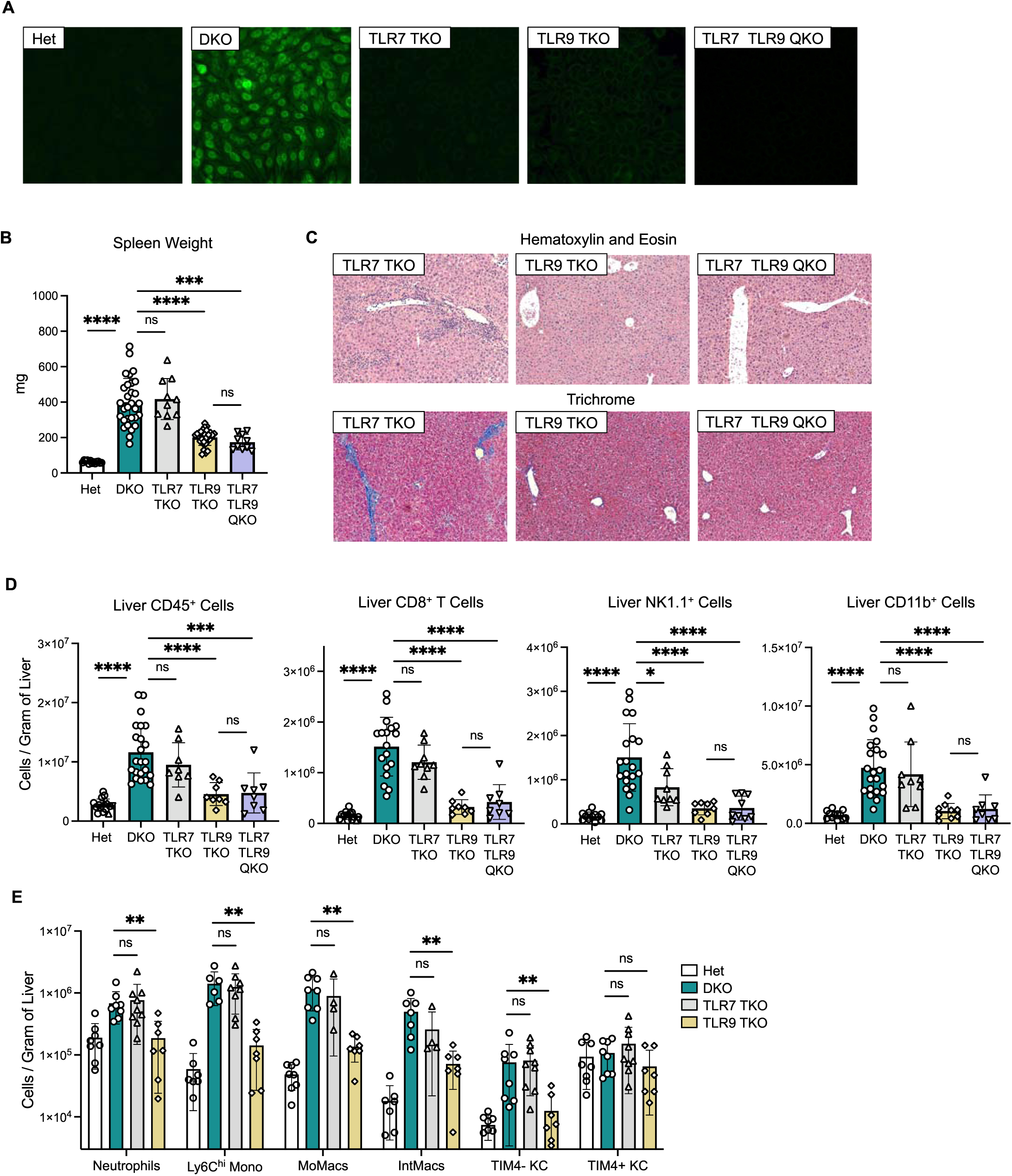
Liver inflammation in DKO mice depends on TLR9. (A) Representative immunofluorescence staining of HEp2 cells with sera from 3-4-month-old mice. (B) Spleen weight of 3-4 month-old mice (n = 10-30 per group). (C) Representative H&E and Masson’s trichrome staining (10x) of liver sections from 3-4 month-old mice. (D) Absolute numbers of the indicated types of immune cells in the livers of 3-4 month-old mice (n = 8-23 per group). (E) Absolute numbers of myeloid subsets in the livers of 3-4 month-old mice (n = 4-9 per group). Data are pooled from at least 5 independent experiments and represented as mean ± SEM. *p < 0.05, **p < 0.01, ***p < 0.001, and ****p < 0.0001. See also Figure S3.

A limited number of patients with mutations in the cytosolic DNA sensor TREX1 have been reported to develop SLE or chilblain-like SLE, presumably through a STING-dependent mechanism (32, 33). However, STING-deficient DKO mice still develop splenomegaly (11, 34), even though they do not develop the inflammatory arthritis that routinely develops in older DKO mice (11). Therefore, it was of interest to determine whether STING contributed to hepatic inflammation in DKO mice. STING^-/-^ DNase1^-/-^ livers were kindly provided by Dr. J. Chen, and by both histologic and flow cytometric criteria, we found that they still developed hepatitis (**Supplemental Figure 3A,B**). Therefore, it appeared that neither STING nor RNA-sensing TLRs contributed to the development of liver disease in DKO mice. Our focus on endosomal TLRs was based on a cross between DKO mice and the ENU loss-of-function mutation Unc93B1^3d^ mice (35), however the Unc93B1^3d^ line had subsequently been shown to have potential off-target effects (14, 36), and so to confirm the importance of endosomal TLRs, we regenerated Unc93B1 TKO mice using an Unc93B1 knockout line produced at UMass Chan by Dr. E. Latz. Again, the Unc93B1 TKO mice failed to develop hepatitis (**Supplemental Figure 3A,B**).

Intriguingly, TLR9 has recently been implicated in several examples of alcoholic and non-alcoholic driven liver inflammation (37–40). Therefore, despite the reported requirement for DNase II in TLR9 activation (19, 41), we decided to explore the role of TLR9 in DKO hepatitis and systemic inflammation and bred TLR9 TKO mice. Unexpectedly, we found that the loss of TLR9 did have a major impact on the extent of autoimmunity and autoinflammation in DKO mice. The TLR9 TKO mice did not make detectable ANAs, presumably because less cell debris accumulated in the absence of inflammation, and splenomegaly was significantly reduced (**Fig. 3A,B**). Liver histology was also dramatically improved (**Figure 3C**). To further assess the impact of TLR9 on liver inflammation, liver cell suspensions were characterized by flow cytometry. As predicted by the histology, the total number of CD45^+^ hematopoietic cells, CD8^+^ T cells, ILCs, and CD11b^+^ myeloid cell were all dramatically reduced but not quite down to the cell numbers of the Het controls (**Figure 3D**). The same trend was seen in the inflammatory myeloid lineage subsets (**Figure 3E**). These parameters were not further reduced in DKO mice deficient for both TLR7 and TLR9 (TLR7 TLR9 QKO). Overall, these data demonstrate a major role for TLR9 in the development of both systemic and liver inflammation in DKO mice, despite the previous studies implicating DNAseII-mediated DNA degradation in the generation of TLR9 ligands. The data further suggest that TLR9 expression in cell types other than B cells or pDCs promotes liver inflammation in DKO mice.

### TLR9 expression in hematopoietic cells is required for liver inflammation

To better understand how TLR9 drives liver inflammation in DKO mice, we used radiation chimeras to ask whether the relevant hepatitis-promoting TLR9-expressing cells were hematopoietic and/or non-hematopoietic cells by generating radiation chimeras in which lethally irradiated DKO or TLR9 TKO were reconstituted with either DKO or TLR9 TKO stem cells. In all cases, the donor and recipient strains differed as far as CD45 alleles such that donor-derived and host-derived cells could be distinguished from one another. These studies included DKOàDKO and HetàHet control groups. The DKOàDKO positive control group fully recapitulated the DKO phenotype (**Figures 1F,4A,B**) and established that a critical disease promoting cell type was not lost by the chimera process.

**Figure 4.**
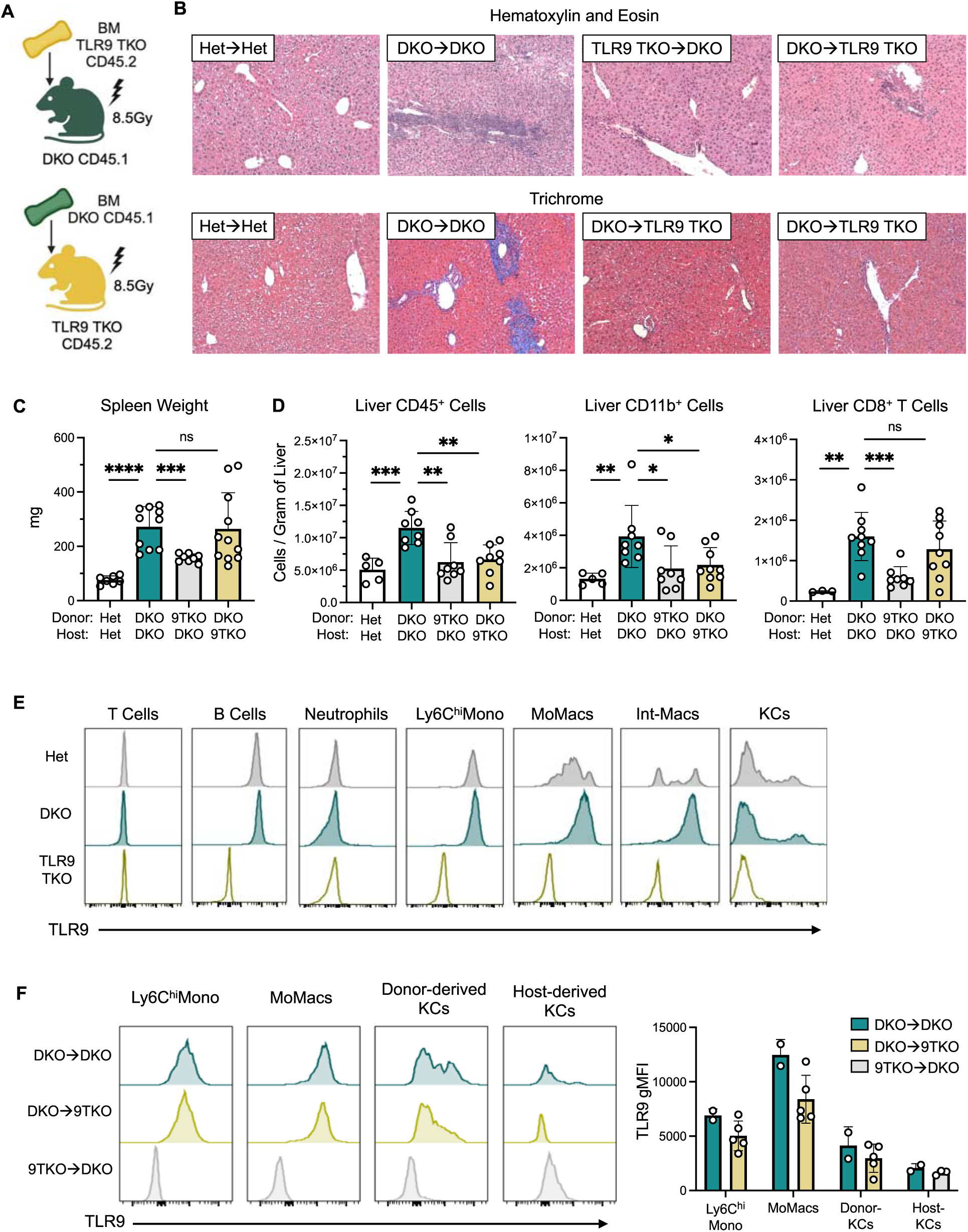
TLR9 in both monocyte lineage cells and liver stromal cells contributes to liver disease in DKO mice. (A) Generating bone marrow chimeras using age- and sex-matched TLR9 TKO C45.2 and DKO CD45.1 mice. (B) Representative H&E and Masson’s trichrome staining (10x) of liver sections from mice 8-week post bone marrow transplant. (C) Spleen weight of mice mice 8-week post bone marrow transplant (n = 7-11 per group). (D) Absolute numbers of indicated immune cells in the livers of mice 8-week post bone marrow transplant (n = 5-9 per group). (E) Representative histograms showing TLR9 expression in each liver immune subset in 3-4-month-old mice. (F) Representative histograms and quantification of TLR9 expression in the indicated myeloid subsets in livers of mice 8 weeks post bone marrow transplant (n = 2-5 per group). Data are pooled from at least 2 independent experiments and represented as mean ± SEM. *p < 0.05, **p < 0.01, ***p < 0.001, and ****p < 0.0001.

We observed that the TLR9 TKOàDKO mice consistently failed to develop hepatitis (**Figure 4A-D**), indicating that a TLR9-expressing bone marrow-derived cell was required for liver inflammation in DKO mice. To further identify the specific bone marrow-derived cell type that contributes to the disease, we first evaluated TLR9 expression in liver immune cells by flow cytometry, comparing to TLR9 TKO mice as negative controls. Intracellular staining for TLR9 showed that neither T cells nor neutrophils expressed TLR9 as expected. We also confirmed TLR9 expression in B cells and Ly6C^hi^ inflammatory monocytes (**Figure 4E**). However, within the macrophage subsets, TLR9 was mostly highly expressed in monocyte-lineage cells including MoMacs and Int-Macs, but not KCs in either Het or DKO mice (**Figure 4E**). We further showed in the chimera mice that the donor-derived newly differentiated KC cells express TLR9 but to a lesser extent than their monocytes and MoMacs precursors, while host-derived radioresistant KCs barely express TLR9 (**Figure 4F**). This data is consistent with previous studies from Barton and colleagues showing downregulation of TLR9 expression in professional scavenger cells (42). Despite TLR9 expression in B cells, it is not likely that TLR9 in B cells is contributing liver disease pathogenesis (19). Instead, our data point to a major role for TLR9-expressing myeloid lineage cells, and in particular MoMacs, in FasL-mediated liver damage.

TLR9 is also expressed in many stromal populations including sinusoidal endothelial cells and hepatocyte. To address the role of TLR9 in the stromal population, we assessed the severity of liver inflammation in DKOàTLR9 TKO mice. Similar to TLR9 TKOàDKO mice, DKOàTLR9 TKO mice showed reduced immune infiltration and fibrosis (**Figure 4B,D**). Since the majority of TLR9 expressing immune cells are bone marrow-derived and we did not detect high level TLR9 expression in radioresistant KCs in either DKOàDKO mice or TLR9 TKOàDKO mice, our data suggests that TLR9 in stromal cells also contributes to the liver inflammation, especially recruitment of the myeloid population.

### TLR9 activated cells drive an amplification loop; IFNγ is required to initiate disease

Despite the remarkable impact of TLR9-deficiency on the development of hepatitis the number of inflammatory monocytes, monocyte-derived macrophages and intermediate monocytes still made up a high proportion of the myeloid lineage cells in the liver of TLR9 TKO mice than in the Het mice (**Figure 5E**), and thus TLR9 TKO mice were not back to baseline. One interpretation of this data was that TLR9 might played more of a role in the amplification of liver inflammation than in the initiation of inflammation.

**Figure 5.**
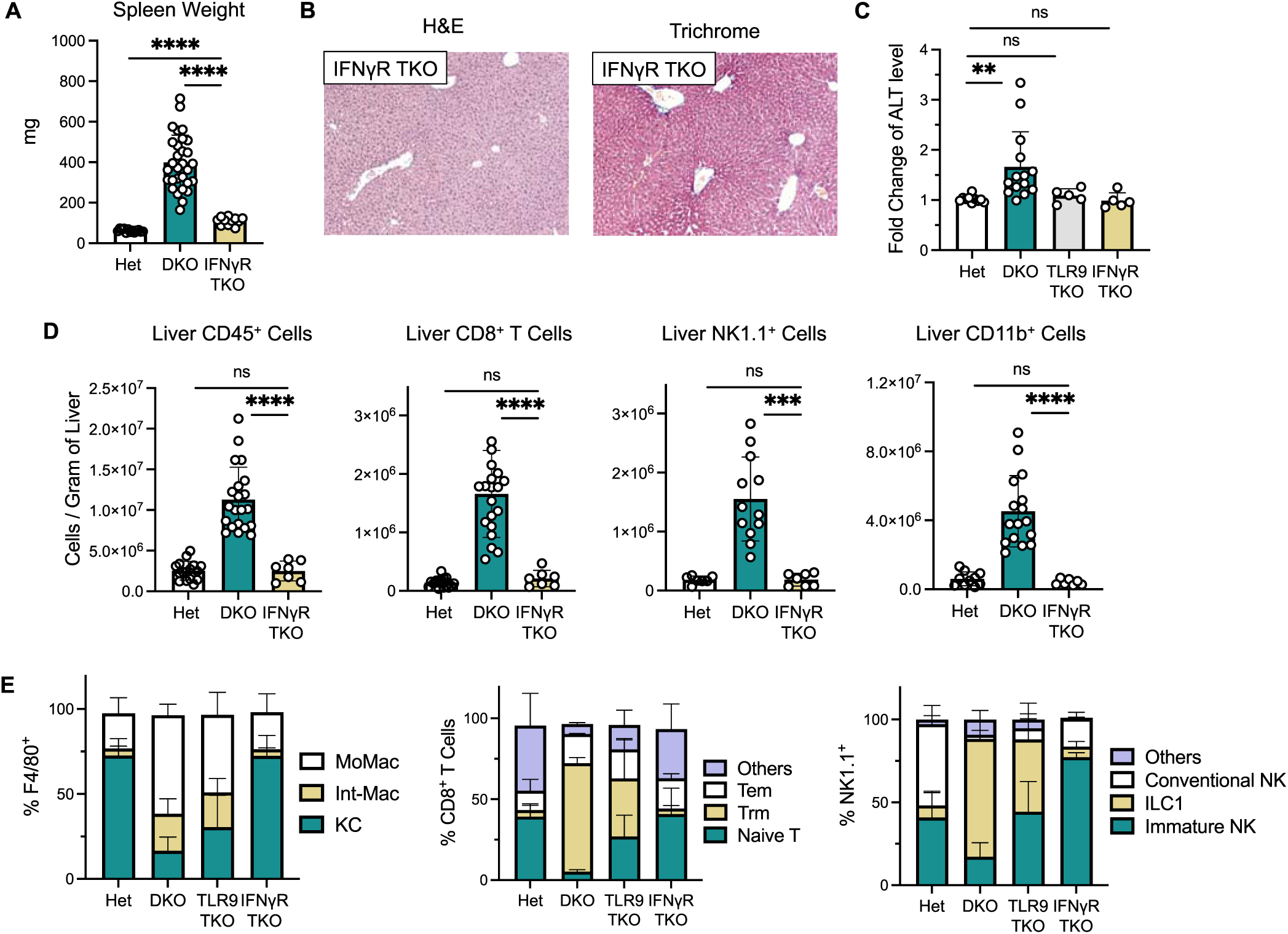
IFNg signaling is required for initiation of immune infiltration in the livers of DKO mice. (A) Spleen weight of 3-4 month-old mice (n = 11-30 per group). (B) Representative H&E and Masson’s trichrome staining (10x) of liver sections from 3-4 month-old mice. (C) Relative ALT level in the sera of 3-4 month-old mice normalized to Het controls (n = 5-18 per group). (D) Absolute numbers of the indicated immune cells in the livers of 3-4-month-old mice (n = 7-21 per group). (E) Frequency of macrophage subsets within liver F4/80^+^ cells, T cell subsets within liver CD8^+^ T cells, and innate lymphocytes cell subsets within liver NK1.1^+^ cells in 3-4 month-old mice. Data are pooled from at least 3 independent experiments and represented as mean ± SEM. *p < 0.05, **p < 0.01, ***p < 0.001, and ****p < 0.0001. See also Figure S4.

One potential upstream signal for the inflammatory process was IFN*γ*. We previously reported an IFN*γ* signature in DKO splenic CD4^+^ T cells (14), and also found a high frequency of IFN*γ*-producing T cells in DKO liver (**Figure 2F**). IFN*γ* has long been recognized as a potent activator of macrophages (6). To further assess the role of IFN*γ* in DKO hepatitis, we intercrossed DKO and IFN*γ*R^-/-^ mice. Remarkably, these IFN*γ*R TKO mice showed no histological evidence of liver inflammation, had fewer liver infiltrating CD45^+^ cells than the TLR9 TKO mice, and also had lower spleen weights (**Figure 5A,B,D**). ALT levels in both the TLR9 TKO and the IFNaR TKO mice were essentially dropped to baseline (**Figure 5C)**. In addition, the number of CD8^+^ T cells, ILC1 cells, and inflammatory myeloid cells was reduced to the same number as the Het controls (**Figure 5D**), Most importantly, in contrast to the TLR9 TKO mice, the frequency of the DKO-amplified subsets essentially returned to the Het ratios (**Figure 5E**). These data suggest that elevated IFN*γ* production precedes the TLR9-driven amplification of liver inflammation in DKO mice. Previously we showed both CD8^+^ and CD4^+^ T cells are actively producing IFN*γ* in the livers of DKO mice (**Figure 2D, Supplemental Figure 2C**), we thus further crossed DKO mice to T and B lymphocyte deficient RAG KO and expect the RAG TKO mice to phenocopy the rescue effect we observed in the livers of IFN*γ*R TKO mice. However, despite improved liver histology, RAG TKO mice still showed increased number of immune cells in the liver featured by monocytic infiltration (**Supplemental Figure 4A,B**). These data suggest the existence of other IFN*γ* producing cells besides T cells that initiate the recruitment of TLR9 expressing macrophages.

## DISCUSSION

Mice deficient for endolysosomal DNase, DNase2 and the type I IFN receptor (IFNaR) develop an autoinflammatory disease that is well-recognized as a model of inflammatory arthritis (8). However, as we have previously described, these mice also develop an SLE-like phenotype associated with splenomegaly, elevated serum cytokines, lymphopenia and autoantibodies (14). In addition, as now described in the current study, similar to patients with hypomorphic mutations in DNAse II, DKO mice spontaneously develop liver disease that recapitulates many of the clinical manifestations of individuals with autoimmune hepatitis (AIH). We now show that inflammation in the DKO liver is characterized by impaired KC resiliency and substantial recruitment of inflammatory myeloid subsets. Concomitantly, populations of chronically activated and/or exhausted lymphocytes, including IFNy producing T cells as well as FasL expressing ILC1s. also accumulate in the liver. The development of AIH depends on both TLR9, and to a greater extent, on IFN*γ*. Additionally, T and B lymphocytes are not required for the recruitment of inflammatory myeloid cells. These findings support a multi-phase positive feedback loop in Dnase2 deficiency autoinflammatory disease wherein IFN*γ* production, by a population of ILCs and/or stromal cells, works synergistically with TLR9 expressing myeloid cells to elaborate fulminate inflammatory liver disease.

AIH is a debilitating disease that affects ∼27/100,00 Americans (18, 43), many of whom do not respond to current therapeutic options. The design of more effective therapies is hampered by the lack of a suitable animal model that mimics the chronic condition found in patient populations. Concanavalin A induced hepatitis, the most commonly studied animal model, depends on the ability of ConA to bind both hepatocytes and the T cell receptor, leading to hepatocyte death and thus elevated serum levels of ALT. However, in contrast to DKO mice, ConA treated mice do not develop liver fibrosis, and depending on the dose, they ConA treated mice either die or the inflammation resolves (15, 16). Other models involve immunization with liver antigens. Again, these models result in transitory inflammation, and fail to capture a chronic progression from early liver immune infiltrate to terminal fibrotic stage (15, 16). By comparison, DKO mice show evidence of portal and interstitial inflammation and fibrosis from a very early age and while most of our data has been obtained from 3-4 month old mice, we have found that even mice that are as young as 4-week-old show portal inflammation, albeit in the absence of fibrosis. Nevertheless, these mice can survive for an extended time period, despite clear histological evidence of liver disease, indicating that important immune regulatory mechanisms are also in play. Therefore, DKO mice provide a unique opportunity to explore the factors that promote chronic inflammation and fibrosis as well as limit the overall extent of liver damage.

The hematopoietic cell types isolated from the livers of DKO mice differ markedly from the hematopoietic cells present in Het control mice and include dramatically expanded numbers CD8^+^ T cells. The DKO CD8^+^ T cells primarily consist of tissue resident memory cells, many of which express PD-1, indicative of some level of T cell exhaustion. Exhausted CD8^+^ T cells have mainly been studied in the context of chronic viral infection or immunosuppressive tumor microenvironments where they are thought to be persistently activated by viral or tumor-associated antigens (44, 45). Establishment of CD8^+^ Trm cells in the liver requires constant antigen stimulation, therefore it is likely that these CD8^+^ T cells in DKO livers are also chronically activated through their TCRs, although their relative affinity for self-antigen may be lower than a typical anti-viral response due to natural tolerance mechanisms that should have limited the development of high affinity self-reactive cells (PMID:30282039). Alternatively, they may recognize antigenic modification that result from ongoing liver inflammation (46). Exhausted T cells have also been identified in the kidneys of SLE-prone MRL/lpr mice, where they retain only a minimal capacity for cytokine production (47). We have found that a remarkably high percentage of the T cells in DKO mice produce, or at least transcribe, IFN*γ*, although they do not appear to express FasL, a proapoptotic molecule commonly produced Tc1 CD8^+^ cytotoxic T cells (48). Exactly how these infiltrating CD8^+^ T cells contribute to the pathogenesis and/or regulation of AIH will be an important topic for future investigations.

DKO livers, and not Het livers, also contain a remarkably high number of ILC1 cells. ILC1s are commonly considered a tissue resident population, primarily known for their ability to produce high levels of IFN*γ*, TNF, and GM-CSF, as well as chemokines involved in the recruitment of other hematopoietic cells to sites of inflammation (24, 49).In addition, it is known that IL15 can induce ILC1 cells to proliferate in the liver and acquire cytotoxic activity (50). In fact, we have found that DKO liver ILC1 cells express incredibly high levels of FasL, even in the absence of metalloproteinase inhibitors, and hepatocytes and other stromal cells in the liver are known to be exquisitely sensitive to FasL-mediated apoptosis (27, 29, 30, 51). Importantly, similar ILC1 cells have recently been detected in the livers of patients with cirrhotic liver disease(52). Together these data suggest that ILC1 play an important role in driving DKO fibrosis and clearly warrant further study.

The major impact of TLR9-deficiency on DKO liver disease was unexpected. As shown in Figure 3A, DKO mice make ANAs, but the staining pattern in a HeEp-2 immunofluorescent assay was consistent with a TLR7 driven response, and in fact autoantibody production is TLR7-dependent. We have further shown that DKO B cells cannot respond to dsDNA internalized through the BCR (19), despite a robust response of DKO B cells to the CpG-rich ODN 1826; the same dsDNA ligands were potent activators of DNase2-sufficient B cells.

Likewise, Miyake and colleagues reported that DKO pDCs could not respond to dsDNA (41). In elegant structural analysis of the endosomal TLRs, this enigma was explained by the identification of two TLR9 DNA binding sites, both of which were required for full activation of this receptor, and both of which are engaged by ODN 1826. One of these two DNA binding sites was found to bind small 5’ DNA fragments and it was proposed that DNase2 likely plays a critical role in the generation of these 5’ fragment (53, 54). Therefore, we assumed that cell debris, resulting from the death of oversaturated DKO scavenger cells, provided the ligands that activated autoreactive DKO B cells specific for RNA-binding proteins, since autoantibody production requires BCR/TLR co-engagement and TLR9 could not be activate d by natural dsDNA ligands in DKO mice. We have now found that TLR9 TKO mice also fail to make ANAs, presumably because there is less inflammation and therefore less debris available for TLR7 B cell activation. To better understand how TLR9 could be promoting AIH, it was important to determine which cells in the DKO liver actually express TLR9.

Intracellular staining revealed that, in addition to B cells, both inflammatory monocytes and monocyte-derived macrophages expressed high levels of TLR9 and are likely to be promoting liver inflammation through a TLR9-dependent mechanism. KCs did not express high levels of TLR9. Myeloid cell activation by TLR9 is likely to depend on a DNAse that is uniquely expressed in cells other than B cells or pDCs. One possibility in DNase1L1 (55). We are in the process of generating DNase1L1 TKO mice to test this hypothesis. However, we cannot rule out a role for TLR9 expression in stromal cells.

The other unanticipated outcome of the current study was the ability of IFN*γ*R-deficiency to completely prevent the development AIH. A number of monogenic autoinflammatory interferonopathies have now been described, most of which have a strong IFN signature and most of which are attributed to type I IFNs (56, 57). However, STING GOF mice that recapitulate the phenotype of SAVI (STING-Associated Vasculopathy of Infancy) patients develop an inflammatory lung disease that depends on IFN*γ* and not type 1 IFNs (58, 59). In fact, IFNaR-deficiency in this model appears to result in more severe disease, pointing to a type I IFN in the negative regulation of inflammation (58, 60). IFN*γ* was originally identified as macrophage activating factor and is now known to sensitize myeloid cells to both TNF and TLR ligands through epigenetic modifications (6, 61). In the context of DKO AIH, both ILC1 cells and CD8 T cells are likely to be important sources of IFN*γ*. Thus we propose that the accumulation of undegraded DNA in liver scavenger cells is likely to be initial event in livers of DKO mice. It is likely that these cells die from necrosis, or another form of inflammatory cell death, as a result of the excessive accumulation DNA. Scavenger cells in the liver include KCs, but also non-hematopoietic cells such as sinusoidal endothelial cells (42), and failure to clear DNA then most likely leads to the production of factors that activate ILC1 cells, leading to their activation and production of IFN*γ*. ILC1 cells further promote the activation and recruitment of myeloid lineage cells and T cells leading to effector populations that further promote inflammation, the activation of hepatic stellate cells and subsequent fibrosis. Admittedly, the details of this process at this point are somewhat vague, but the similarities of this model to the AIH described in patient populations makes a strong case for further mechanistic studies. The current report also identifies several key therapeutic targets including TLR9 and type II interferons.

## MATERIALS AND METHODS

### Mice

DNase II-gene targeted mice were kindly provided by Dr. S. Nagata and obtained through the Riken Institute. Dnase2+/− x Ifnar−/− (Het), Dnase2−/− x Ifnar−/− (DKO), Dnase2−/− x Ifnar−/− x Tlr7−/− (TLR7 TKO), Dnase2−/− x Ifnar−/− x Ifngr−/− (IFNGR TKO) mice have been described previously (14). Dnase2*^−/−^* Sting^Gt/Gt^ mice were kindly provided by Dr. Z. Chen (UT Southwestern Medical Center, Dallas, TX). Tlr9^-/-^ (Jax #034449), IFN*γ* reporter B6.129S4-Ifng^tm3.1Lky^/J (Jax# 017581), and B6 CD45.1 (Jax #002014) mice were obtained from Jackson Laboratory. Unc93b1^-/-^ mice were kindly provided by Dr. E. Latz. DKO mice were intercrossed with Tlr9^-/-^, Unc93b1^-/-^, B6.129S4-Ifng^tm3.1Lky^/J, and B6 CD45.1 mice to generate the corresponding triple KO lines, DKO X GREAT line, and CD45 allotype distinct line. TLR9 TKO mice were further intercrossed with TLR7 TKO mice to generate TLR7 TLR9 QKO mice. Mice were euthanized by isoflurane. Serum was collected by cardiac puncture on euthanized animals. All animal procedures were proved and performed in accordance *with the* Institutional Animal Care and Use Committee at the University of Massachusetts Medical School.

### Histological Analysis

The left lateral lobe of livers was removed, fixed with 10% phosphate buffered formalin (PBF) at room temperature, and embedded in paraffin. Hematoxylin and eosin (H&E), Masson’s trichrome, and TUNEL staining on paraffin-embedded liver sections were carried out by the Morphology Core (University of Massachusetts Chan Medical School, Worcester, MA). Samples were observed under EVOS M7000 microscope in the Bone Analysis Core (University of Massachusetts Chan Medical School, Worcester, MA).

### Plasma alanine aminotransferase activity

Serum samples were collected from 3–4-month-old mice. Serum levels of alanine aminotransferase (ALT) activity were determined using the Alanine Transaminase Activity Assay Kit (Abcam #ab105134) following manufacturer’s instruction.

### Liver processing and cell suspension

To assess immune cells from the liver by flow cytometry, livers were perfused with 10 ml ice-cold PBS through the portal vein, then homogenized using Liver Digestion Kit (Miltenyi #130-105-807) and GentleMACS Octo Dissociator (Miltenyi). Cell suspensions were passed through a 100uM cell strainer, washed with plain DMEM and PBS containing 2mM EDTA and 0.5% BSA, and centrifuged at 300g for 10 min. RBCs were removed with Hybri-Max RBC lysing buffer (Sigma) before staining.

### Flow cytometry and cell sorting

Cell suspensions were incubated with anti 2.4G2 antibody to block Fc receptors and stained with fluorophore conjugated primary antibodies at room temperature for 25-30 min. Samples were fixed using FluoroFix buffer (Biolegend #422101). TLR9 intracellular staining was performed on extracellularly stained cells using the Fixation/Permeabilization Solution kit (BD Biosciences #554714) following the manufacturer’s protocol. Ki-67 nuclear staining was performed on extracellularly stained cells using the FoxP3/Transcription factor staining buffer kit (Invitrogen #00-5523-00) according to manufacturer’s protocol. The absolute cell counts were determined using Precision Count Beads (Biolegend #424902) and expressed as the number of cells per gram of tissue. Data were acquired on a 5-laser Aurora (Cytek) and analyzed with FlowJo software (Tree Star).

Antibodies were purchased from BioLegend, ThermoFisher, Tonbo Biosciences, eBioscience, and BD Biosciences Pharmingen. The following primary antibodies were used for staining:

CD45 BV650 (30-F11), CD11b BUV805 (M1/70), Ly6G BV421 (1A8), MHCII Pacific Blue (M5/114.15.2), CD11c BV570 (N418), CD86 BV785 (GL-1), Ly6C FITC (HK1.4), TIM4 PE (F31-5G3), CD3 PE-Cy5 (17A2), F4/80 APC-Cy7 (BM8), CLEC2 APC (17D9), CD19 APC-Fire810 (6D5), MHCI AF700 (AF6-88.5), CD8 BUV395 (53-6.7), TCRb BUV737 (H57-597), CD44 BV605 (IM7), DX5 FITC (DX5), NK1.1 PE (PK136), CXCR6 PE-Dazzle594 (SA051D1), CD49a PE-Cy7 (HMa1), CD4 SparkNIR685 (RM4-4), CD62L APC-efluor780 (MEL-14), PD-1 APC-Fire810 (29F.1A12), FasL APC (MLF3), TLR9 PE (S18025A), TLR9 APC (S18025A), Ki-67 FITC (11F16) CD45.1 AF700 (A20), CD45.2 BV750 (104). All cells were stained for viability with Ghost Violet510 (Tonbo).

### Antinuclear antibody assay

Anti-nuclear antibody assay was performed as described previously (14). In brief, slides coated with Hep-2 cells were stained with serum samples (1:50 dilution) collected from 3–4-month-old mice. Bound antibodies were detected with DyLight 488-coupled goat anti-mouse IgG Ab (Poly4053, BioLegend). Samples were visualized by fluorescent microscopy.

### Generation of Radiation Chimeras

Lethally irradiated (8.5Gy) 7-9-week-old mice were reconstituted with 10^7^ bone marrow cells from sex and age matched mice by i.v. injection. Mice were then maintained on sulfatrim water. Chimerism and disease severity were assessed 9 weeks post reconstitution. The extent of reconstitution was determined by flow cytometry based on CD45 allelic marker.

### Statistical Analysis

All statistical analyses were performed in Prism V.10 (GraphPad Software). Sample size in each experiment is detailed in the figure legend, where n = number of mice. Normality of the data distribution was examined by Shapiro-Wilk tests. The statistical significance of differences between two groups was determined by student t-test for normally distributed data, and by nonparametric tests for data that are non-normal distributed. One-way ANOVA was used when comparing more than two groups. Differences were considered significant if P < 0.05, and the following represents the level of significance: ****P < 0.0001, ***P < 0.001, **P < 0.01, and *P < 0.05.

## ACKNOWLEDGEMENT

We would like to thank Sruthi Takillapati and Stephanie Moses for genotyping and maintaining the mouse colony. Thank you to our colleagues Drs. Kerstin Nundel, Michelle Kelliher, Pranoti Mandrekar and Kate Fitzgerald constructive feedback and discussion. We also thank Dr. Zhijian Chen for providing tissues from Dnase2*^−/−^* Sting^Gt/Gt^ mice. We would like to thank Dr. Marc Sherman (MGH) for assistance with evaluation of liver sections. These studies were supported by NIH grant R37 AI155901.

---

**Figure S1.**
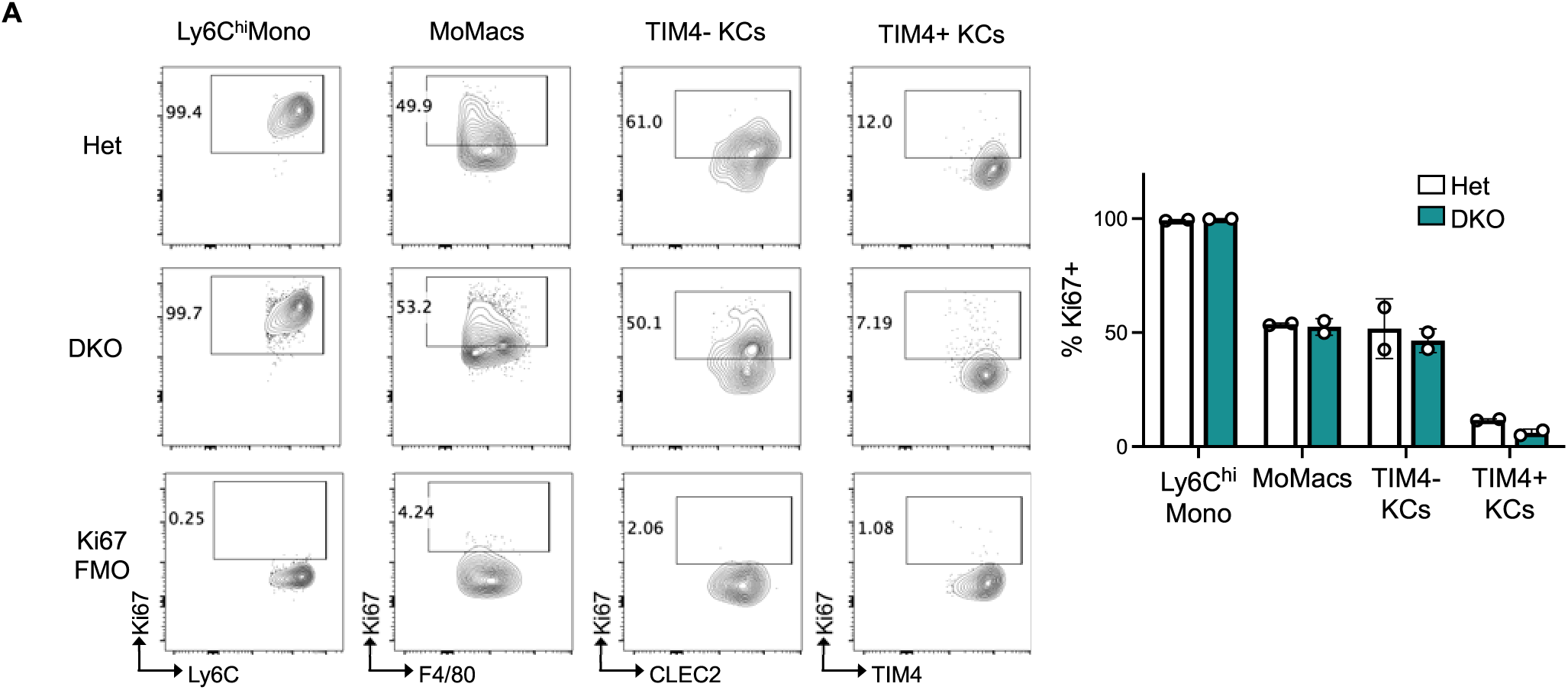
Monocytes lose proliferative potentials as they differentiate into Kupffer cells. (A) Representative flow plots and quantification of Ki-67^+^ cells among liver monocytes, MoMacs, TIM4^+^ KCs, and TIM4^-^ KCs in 3-4-month-old mice (n = 2 per group). Data are represented as mean ± SEM.

**Figure S2.**
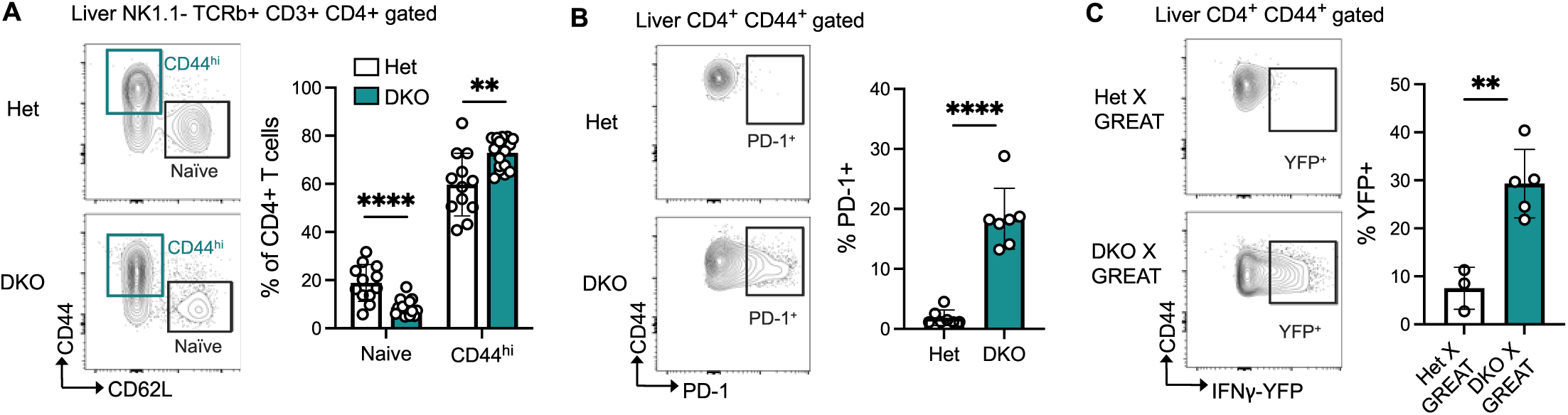
Activated CD4^+^ T cells produce IFNg in the livers of DKO mice. (A) Representative CD44 and CD62L staining of CD4^+^ gated liver T cells. Bar graph compares the frequency of naïve and CD44^hi^ activated CD4^+^ T cells in livers of 3-4 month-old mice (n = 12-17 per group). (B) Representative PD-1 staining of CD44^+^ gated liver CD4^+^ T cells of 3-4-month-old mice. Bar graph compares the frequency of PD-1^+^ cells within CD44^hi^ CD4^+^ T cells in the liver (n = 7 per group). (C) Representative YFP flow plot of CD44^+^ gated liver CD4^+^ T cells of 3-4-month-old IFNγ reporter mice. Bar graph compares the frequency of YFP^+^ cells within CD44^hi^ CD4^+^ T cells in the liver (n = 3-5 per group). Data are represented as mean ± SEM. *p < 0.05, **p < 0.01, ***p < 0.001, and ****p < 0.0001.

**Figure S3.**
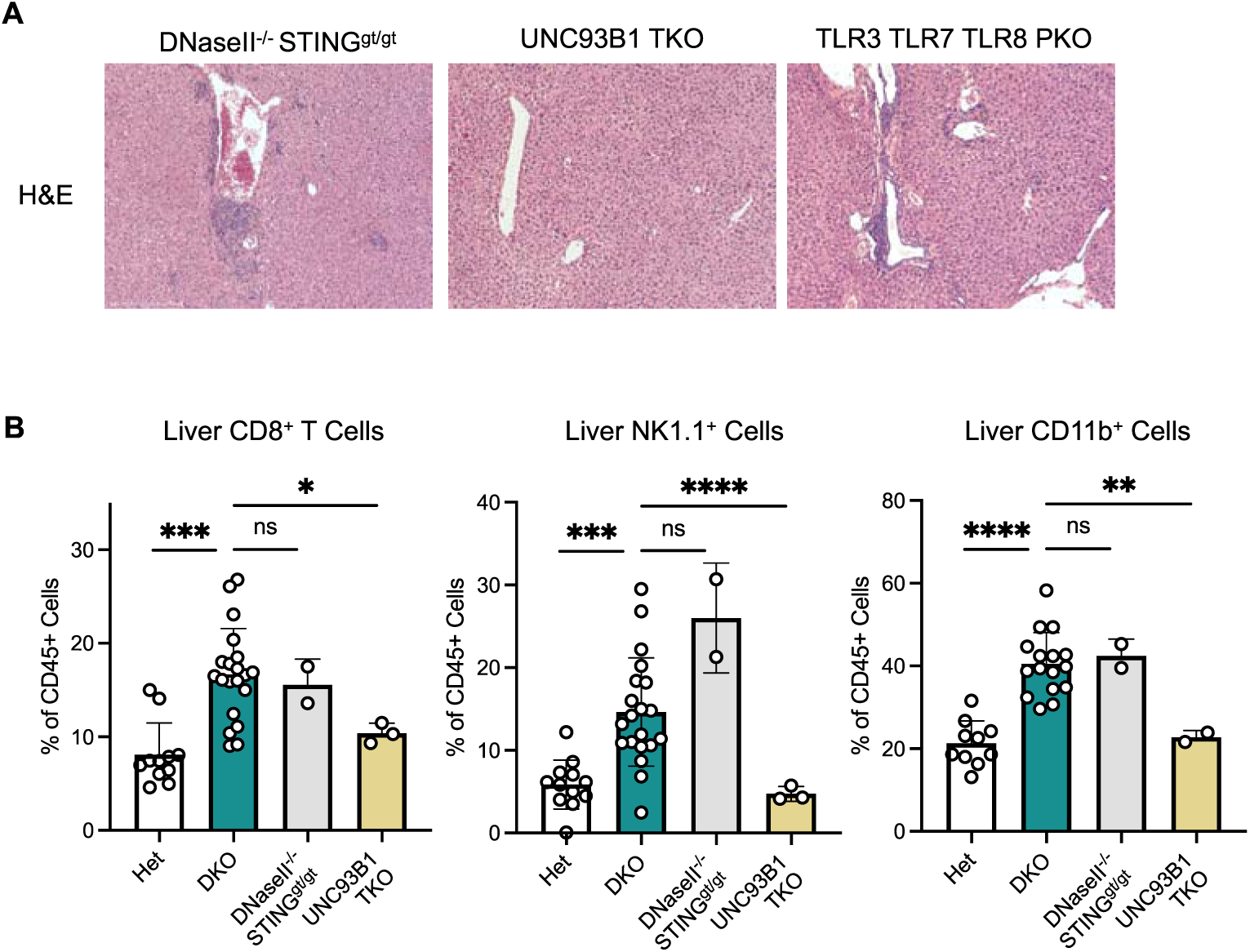
Liver immune infiltration in DKO mice depends on endosomal TLRs but not STING. (A) Representative H&E staining (10x) of liver sections from 3-4 month-old mice. (B) Frequency of indicated immune population within liver CD45^+^ cells in 3-4 month-old mice (n = 2-15 per group). Data are represented as mean ± SEM. *p < 0.05, **p < 0.01, ***p < 0.001, and ****p < 0.0001.

**Figure S4.**
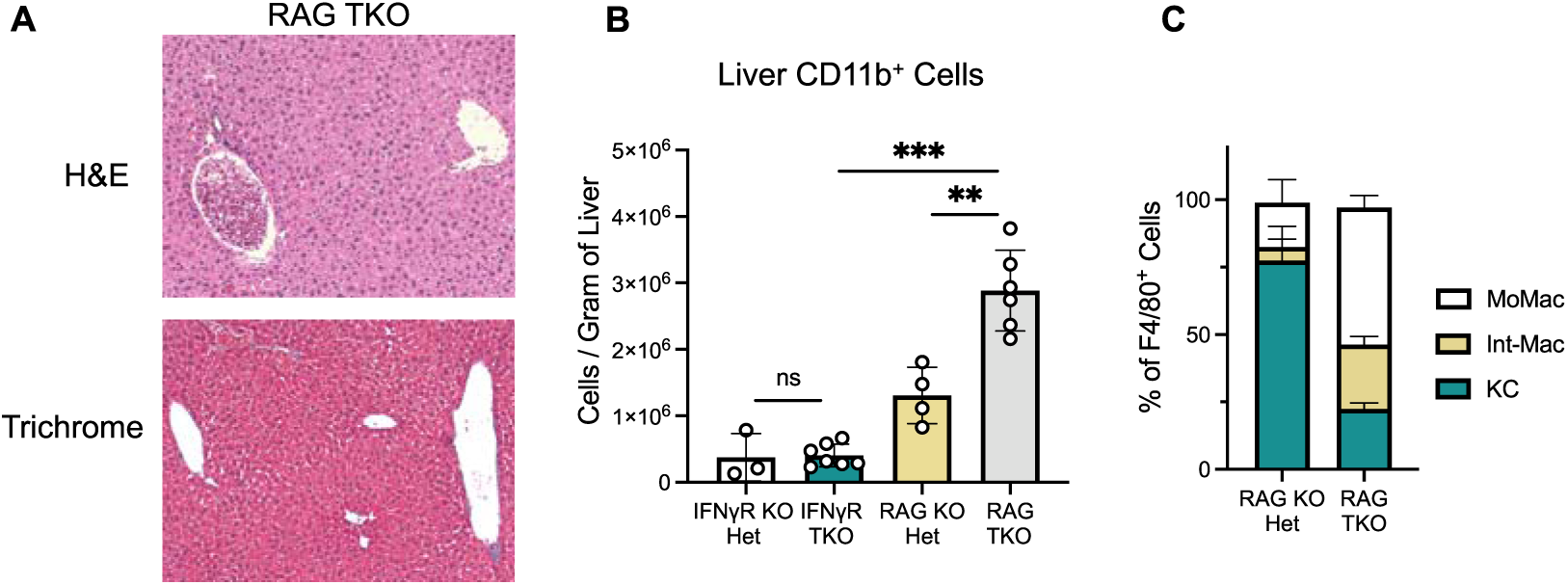
Increased number of myeloid cells in the livers of RAG TKO mice. (A) Representative H&E and trichrome staining (10x) of liver sections from 3-4 month-old mice. (B) Absolute number of CD11b^+^ myeloid cells in the livers of 3-4 month-old mice. (C) Frequency of macrophages subsets within liver F4/80^+^ cells in 3-4 month-old mice. Data are represented as mean ± SEM. *p < 0.05, **p < 0.01, ***p < 0.001, and ****p < 0.0001.

